# First evidence of a menstruating rodent: the spiny mouse (*Acomys cahirinus*)

**DOI:** 10.1101/056895

**Authors:** N. Bellofiore, S. Ellery, J. Mamrot, D. Walker, P. Temple-Smith, H. Dickinson

## Abstract

**Background:** Advances in research relating to menstruation and associated disorders (such as endometriosis and pre-menstrual syndrome) have been hindered by the lack of an appropriate animal model. Thus, many aspects of this phenomenon remain poorly understood limiting the development of efficacious treatment for women. Menstruating species account for only 1.5% of mammals, and less than 0.09% of these are non-primates. Menstruation occurs as a consequence of progesterone priming of the endometrial stroma and a spontaneous decidual reaction. At the end of each infertile cycle as progesterone levels decline the uterus is unable to maintain this terminally differentiated stroma and the superficial endometrium is shed. True menstruation has never been reported in rodents.

**Objective:** Here we describe the first observation of menstruation in a rodent, the spiny mouse (*Acomys cahirinus*).

**Study Design:** Virgin female spiny mice (n=14) aged 12-16 weeks were sampled through daily vaginal lavage for 2 complete reproductive cycles in our in-house colony at Monash Medical Centre, Clayton, Australia. Stage-specific collection of reproductive tissue and plasma was used for histology, prolactin immunohistochemistry, and ELISA assay of progesterone (n=5 / stage of the menstrual cycle). Normally distributed data are reported as the mean ± standard error and significant differences calculated using a one-way ANOVA. Non-normal data are displayed as the median values of replicates (with interquartile range) and significant differences calculated using Kruskal-Wallis test.

**Results:** Mean cycle length was 8.7 ± 0.4 days with red blood cells observed in the lavages over 3.0 ± 0.2 days. Cyclic endometrial shedding and blood in the vaginal canal concluding with each infertile cycle was confirmed in all virgin females. The endometrium was thickest during the luteal phase, when plasma progesterone peaked at ~102.1 ng/mL and the optical density for prolactin immunoreactivity was strongest. The spiny mouse undergoes spontaneous decidualisation, demonstrating for the first time menstruation in a rodent.

**Conclusion:** The spiny mouse is the first rodent species known to menstruate and provides an unprecedented natural non-primate model to study the mechanisms of menstrual shedding and repair, and may be useful in furthering our understanding of human specific menstrual and pregnancy associated diseases.

## INTRODUCTION

Menstruation, the cyclical shedding of the decidualised endometrium in the absence of pregnancy, is believed to be limited to 78 higher order primates (humans and Old World monkeys), 4 species of bat[1, 2] and the elephant shrew[1, 3, 4]. This represents only 1.51% of the known 5502 mammalian species[5]; <0.09% of menstruating species are nonprimates. Common to these species, and inherent to the process of menstruation, is spontaneous decidualisation of the endometrial stroma without initiation from an implanting embryo. Under the control of progesterone from the ovary, the decidual reaction occurs in unison with a series of intricate structural changes to the uterine stratum functionalis, including extensive angiogenesis of maternal vasculature into spiral arterioles[6]. In the absence of pregnancy, degeneration of the corpus luteum results in progesterone withdrawal and endometrial shedding which, due to extensive vascularization of the endometrium, is accompanied by bleeding into the uterine cavity[1]. In non-menstruating species, decidualisation of the endometrium does not eventuate unless fertilization occurs and the process is signalled from the conceptus.

The spontaneous nature of the morphological changes that result in decidualisation of the endometrial stroma are considered a pre-emptive maternal defence against the invasiveness of the trophoblast[1, 7, 8]. The extent of trophoblastic invasion is severe in menstruating species; and can reach as far as the inner third of the myometrium in women, resulting in what is considered to be an aggressive form of haemochorial placentation[9]. Ultimately for successful pregnancy to occur, the maternal decidual reaction must be precisely balanced to allow adequate blood flow and nutrient supply to the invading trophoblast but to also to ensure the mother is sufficiently protected[1]. Alterations in this delicate balance manifest as clinical diseases such as pre-eclampsia; currently one of the leading causes of fetal-maternal morbidity and mortality. Pre-eclampsia is thought be due to inadequate vascular remodelling and an impaired decidual reaction, resulting in shallow trophoblastic invasion and placental hypoxia [10]. Alternatively, if the trophoblast invades too deeply, women may suffer placenta acreta, with abnormal placental-uterine adhesion. In extreme cases, this may only be able to be treated with peri-partum hysterectomy[11].

Endometriosis, resulting from the presence of endometrial tissue outside the uterine cavity, affects up to 10% of women with symptoms such as dysmenorrhea, chronic pelvic pain, dyspareunia, and infertility. As for pre-eclampsia and placenta acreta, the aetiology of endometriosis is unknown and there is no cure, with treatments only targeting symptoms and not the underlying causes. Although mouse models of artificial menstruation exist[12, 13], the limited research into menstruation and its related disorders is largely due to the absence of a cost-effective and practical laboratory model of natural menstruation. This study describes the first report of menstruation in a rodent: the common or Cairo spiny mouse (*Acomys cahirinus*), which is native to Northern Africa. The spiny mouse produces small litters (typically 2-3) of precocial pups, with most organogenesis completed *in utero* during the relatively long (for rodents) gestation of 39 days[14]. Initial observations of blood at the vaginal opening of some female spiny mice in our breeding colony lead to this unexpected discovery.

## MATERIALS & METHODS

**Animal Care**. All experiments were conducted in accordance with the Australian Code of Practice for the Care and Use of Animals for Scientific Purposes, and all experiments approved by the Monash University/Monash Medical Centre Animal Ethics Committee. Virgin female spiny mice (n=18) aged 12-16 weeks, were housed in groups of 5-6 per cage. Cages were lined in wooden shavings and cardboard enrichment provided in forms of tunnels and climbing apparatuses. Food (Rat and Mouse Cubes, Specialty Feeds, WA) and water were provided *ad libitum* and supplemental fresh vegetables (carrots and celery, up to 50g per cage) were provided weekly. Animals were checked at least twice daily and cages were cleaned weekly. Female weights were monitored daily throughout the experiment. These animals were sourced from our own research colony, where temperature is maintained at 25-27°C, humidity 30-40%, with a 12h light-dark cycle (lights on 07:00)[14].

**Vaginal Lavage**. Vaginal lavages were performed daily between 12:30 and 14:30 using sterile saline solution (0.9% w/v), and the time of sampling recorded each day. Female spiny mice were restrained in the supine position with a small handtowel by gently scruffing behind the neck and shoulders (a handtowel is necessary due to the fragility of spiny mouse connective tissues and the propensity for skin to easily tear). The vulva was then lubricated with non-scented, water-based lubricant (Ansell, VIC). A 1 mL plastic transfer pipette was used to draw up ~50 μL of saline, before gentle insertion into the vaginal canal, no deeper than 8 mm, and well before contact with the cervix. The saline was flushed into the vaginal cavity twice, the solution redrawn into the pipette, the pipette removed and the sample expelled onto a glass histological slides (Menzel-Gläser Superfrost, Germany). Samples were dried at 27°C for 5 min then sprayed with cytology fixative (Spray Fix, Surgipath Medical Industries, USA). Smearing occurred consecutively for 18 days. Cycle stages were distinguished based on the dominant cell type(s) present in the smear[15, 16]. If the smear appeared to be in a transitional phase i.e., between two consecutive stages, each of the stages was designated a time of 0.5 days to the total length. Otherwise, when a particular smear was observed, it was designated a period of 1 day to the total length for that stage.

For technical comparison, F_1_ (C57BL/6J X CBA/J) hybrid mice (n=5) sourced from the Monash Animal Research Platform weighing 17-20g (40-60 days old) were smeared daily for 12 consecutive days. Females were held in the same supine position but in the absence of a towel. All other aspects of smearing technique were the same as described for the spiny mouse above.

**Cytology Staining** Slides were stained with haematoxylin and eosin (H&E) (Amber Scientific; Harris Haematoxylin and 1% aqueous Eosin, WA). Briefly, slides were rehydrated in running tap water for 30 sec −1 min prior to haematoxylin staining for 5-6 minutes. Slides were then washed thrice in tap water to remove excess stain and differentiated in 5% acid ethanol. Following rinsing, slides were submerged repeatedly in a 5% ammonia solution to develop blue coloration. Slides were counterstained with eosin for 3 min. Slides were progressively dehydrated in graded ethanol solutions and underwent three successive changes of clearing xylene for 2 min each. Each slide was cover-slipped with DPX mounting-medium.

**Post Mortem Analysis and Tissue Collection**. Female spiny mice (n=4) were killed humanely at each distinct stage of the menstrual cycle, which were confirmed by vaginal smear prior to post-mortem. Animals were subjected to heavy anaesthetic (95% isoflurane) before cardiac puncture, followed immediately by cervical dislocation. Whole blood (0.3-1.2 mL per animal) was collected in a heparin-lined tube and subjected to plasmapheresis in a refrigerated centrifuge (3000 RPM at 4°C for 10 min). The separated plasma was then frozen at −20°C for later analysis. The whole, intact uterus with both ovaries and the cervix attached was trimmed of fat and removed. Ovaries were then separated, and the uterus weighed (wet weight) before fixation in 10% buffered formalin for 48h, followed by 70% ethanol for 24-72h. Samples were processed to paraffin wax using a Leica ASP-300 processer, before being embedded in paraplast paraffin medium. Tissues were sectioned (5 μm), adhered to slides and baked at 60°C for 20-30 min. Samples were dewaxed through successive xylene changes, cleared in graded ethanol and rehydrated in tap water.

**Histological Staining**. To visualise reproductive tissues, slides were stained with Mallory’s trichrome. Briefly, sections underwent a secondary fixation in Bouin’s fluid at 60°C for 60 min prior to dewaxing. Following this, tissues were stained with acid fuschin 1g/100mL dH20 for 2 min, rinsed thoroughly in distilled water and stained with phosphomolybdic acid 1g/100mL dH_2_O for 2 min. Slides were rinsed, before staining with Orange G 2g, Methyl blue 0.5g, oxalic acid 2g/100mL dH_2_O for 15 minutes. Following thorough rinsing, tissues were dehydrated and differentiated as for H&E above.

**Progesterone Assay**. Plasma progesterone concentration was measured using a commercially available mouse/rat enzyme-linked immunosorbent assay kit (ALPCO, #55-PROMS-E01). All samples were measured in duplicate and the median value reported. The validity of spiny mouse plasma levels was tested by performing spike and recovery and linearity of dilution procedures following manufacturer’s protocols. Selected samples from individual animals were spiked with 0, low (10 ng/mL), medium (25 ng/mL) or high (50 ng/mL) analyte (Progesterone powder, Sigma-Aldrich, NSW) and one sample diluted by a factor of 4, 8 and 16, and the expected vs. observed recovery measured.

**Immunohistochemistry**. Fixed tissues were subjected to immunohistochemistry to detect presence of decidualisation using prolactin as biomarker[17, 18]. Samples were dewaxed through multiple changes of histolene, and rehydrated through decreasing concentrations of ethanol until submerged in running tap water for 1 min. Slides were washed in TBS-Tween 20 (0.1%) buffer solution thrice for 5 min each before exogenous peroxidase activity was blocked by applying 1% H_2_O_2_ solution for 20 min. Slides were thrice washed in buffer and serum block was applied for 30 min at room temperature (Dako, X090930-2, NSW). Slides were incubated in primary polyclonal rabbit anti-human prolactin (Dako, A0569, NSW at 1:400 dilution in 10% NGS at 4°C overnight or 10% normal goat serum as a negative control. Slides were thrice washed and incubated in secondary biotinylated goat anti-rabbit IgG (Vector Laboratories, CA) at 1:200 dilution for 60 min at room temperature. Following washing, slides were incubated in avidin-biotin complex at room temperature for 60 min, then washed. 3,3’-Diaminobenzidine was then applied for 6 min for visualisation. Slides were buffer washed thrice and washed again in running tap water before dehydration in decreasing graded ethanol. Slides were cleaned and coverslipped as in the H&E protocol outlined above. Uterine tissue from an early gestational females (7 days gestational age) were used as positive controls for prolactin staining (not shown).

**Image Acquisition**. Images of whole organs were captured using an iPhone 4. All histological images were captured using a Leitz Diaplan imaging bright-field microscope, with light settings kept constant and analysed with the programs Axio-Vision version 4.6.3 and Image Pro-Plus version 6. Images of the vaginal cytology of both the spiny mouse and F1 mouse were taken at X200 magnification. Histological images of the cervix and ovaries of the spiny mouse were taken at X40 magnification, and the sections of the uterine horns taken at X100 and X200 magnification. Using Axio-Vision, the thickness of the endometrium and myometrium were measured, as was the diameter of the uterine lumen (n=12 measurements per stage), across all stages of the reproductive cycle in the spiny mouse (μm). The degree of immunopositive staining was determined using Image Pro-Plus, as described by Atik (2014)[19]. Calibration of the light and dark fields was implemented by the use of a control image containing both incident light and infinite optical density; brought about by use of a scalpel blade placed on the stage to partially obscure the light. All images were captured in one session, with all program settings and light exposure kept exact for each section. Images were converted to grayscale and the mean optical density of positive prolactin-stained areas of the endometrium and myometrium were measured. Mean values were calculated for each section, with 3 fields of view at X200 magnification per structure of interest (endometrium or myometrium) per animal. Regions containing background staining were measured for each image and subtracted from the mean optical density of each sample to adjust for non-specific staining. The mean of means for each group was then calculated. The investigators conducting measurements of the reproductive tract and optical density were blinded to the treatment groups at the time of analysis.

**Image Alterations**. After completion of analysis, images were subjected to minor aesthetic alterations using Adobe Photoshop CC (2014). Images were sharpened, and backgrounds brightened if required. Due to the variability in staining, colour balancing was performed on some histological images to match specific sections of tissues for easier comparison.

**Statistical Analysis**. All statistical analyses were conducted using SPSS (Version 22) and GraphPad Prism (Version 6.01). Normality of the data was tested before statistical analysis. Uterine weight, endometrial and myometrial thickness, uterine lumen diameter and plasma progesterone concentrations (n=5 samples per stage) are displayed as the median values of replicates (with interquartile range). Significant differences were calculated using Kruskal-Wallis. Optical density, menstrual cycle length are reported as the mean ± standard error (SEM). Significant differences (corresponding to a p value of < 0.05) were calculated using a one-way ANOVA. Tukey’s multiple comparisons test for comparing all other parameters between stages of the reproductive cycle during post-hoc analysis.

## RESULTS

We examined cytology of daily vaginal lavages from virgin females (n=14) and found an overall cycle length ranging from 6-10 days, with an average of 8.7 ± 0.4 days. Cytology showed all of the expected stages of a rodent oestrous cycle: a follicular phase comprising proestrus and oestrus and a luteal phase denoted by metestrus and diestrus. Spiny mice have an additional stage not seen in rodents, which is characterised by the presence of large numbers of red blood cells (Figure 1) over a period of 3.0 ± 0.2 days, consistent with menses. Blood was present in the vaginal lavage of all females (14/14) during the transition from the luteal to the follicular phase in both cycles studied. Blood was visible macroscopically on the external genitalia in 4 subjects (29%) during the lavage process.

**Figure 1:**
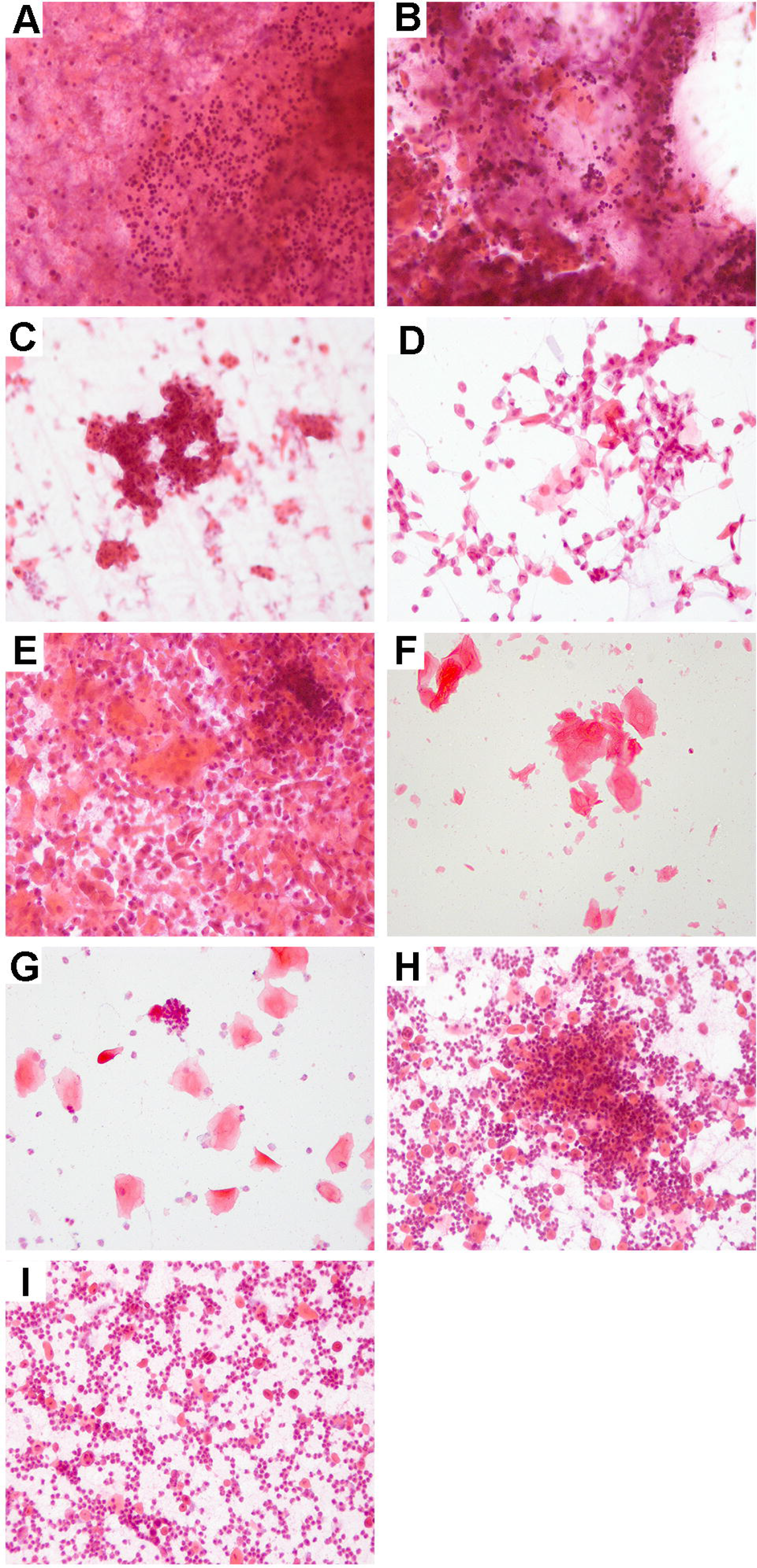
Vaginal cytology of the female spiny mouse. Cytology showing a female spiny mouse with a 9 day cycle. Early menses at the conclusion of the previous infertile cycle (**A, B**). Proestrus, the beginning of the follicular phase; containing nucleated epithelial cells (**C, D**). Transition to oestrus (**E**). Oestrus (**F**), characterised by cornified epithelial cells. Metestrus; transitioning to the luteal phase (**G**). Diestrus, the luteal phase, containing high leukocytic infiltration (**H, I**). Menses will follow within 24-48h. Scale bars = 50 μm, magnification 200X, Haematoxylin and Eosin stain.

Reproductive tract dissections revealed significant differences in uterine weight during the cycle (Table 1), with blood present in the uterine lumen corresponding to the time it was detected in the smears (Figure 2). The timing and recurrence of bleeding was firm evidence for menstruation in the spiny mouse.

**Table 1:** Significant changes during the menstrual cycle of the spiny mouse.

|  | *Follicular Phase* |  | *Luteal Phase* |  | *Menses* |
| --- | --- | --- | --- | --- | --- |
|  | Proestrus | Oestrus | Metestrus | Diestrus |  |
| Length of Cycle Stage (Days)<sup>1</sup> | **1.80** ± 0.2 | **1.30** ± 0.2 | **0.71** ± 0.1 | **1.80** ± 0.3 | **3.00** ± 0.2 |
| Uterine Wet Weight (g)<sup>2</sup> | **0.26**<br>(0.22, 0.29) | **0.32**<br>(0.27, 0.33)<sup>a,b</sup> | **0.15**<br>(0.13, 0.16) | **0.09**<br>(0.07, 0.11)<sup>a</sup> | **0.12**<br>(0.09, 0.16)<sup>b</sup> |
| Endometrial Thickness (µm)<sup>2</sup> | **64.7**<br>(53.49, 144.3)<sup>a</sup> | **112.7**<br>(73.60, 181.4)<sup>b</sup> | **225.9**<br>(180.6, 375.7)<sup>a</sup> | **322.6**<br>(254.8, 512.2)<sup>a,b</sup> | **213.3**<br>(171.3, 300.1)<sup>a</sup> |
| Diameter of Uterine Lumen (µm)<sup>2</sup> | **671.1**<br>(437.9, 1378.0)<sup>a</sup> | **1334.0**<br>(1158.0, 1868.0)<sup>b</sup> | **345.9**<br>(131.6, 772.9)<sup>b</sup> | **127.0**<br>(110.3, 192.3)<sup>a,b</sup> | **614.9**<br>(228.4, 840.1)<sup>b</sup> |
| Endometrial Optical Density (Prolactin)<sup>1</sup> | **0.03** ± 0.01<sup>a</sup> | **0.05** ± 0.01 | **0.05** ± 0.01 | **0.07** ± 0.01<sup>a</sup> | **0.06** ± 0.01 |
| Plasma Progesterone (ng/mL)<sup>2</sup> | **46.6**<br>(29.4, 63.9) | **65.38**<br>(32.9, 82.7) | **42.9**<br>(37.2, 54.6) | **102.1**<br>(70.9, 198.6)<sup>a</sup> | **50.5**<br>(30.6, 65.4)<sup>a</sup> |

Superscript letters correspond to groups significantly different from each other. Data subjected to normality tests. For normally distributed data, the mean ± the standard error is reported<sup>1</sup>. The median (25<sup>th</sup> percentile, 75<sup>th</sup> percentile) is reported for all other values<sup>2</sup>. Significant differences corresponded to a p value of <0.05 (one-way ANOVA and Dunnett's post-hoc for normally distributed data and Kruskal-Wallis and Dunn's post-hoc for all other data).

**Figure 2:**
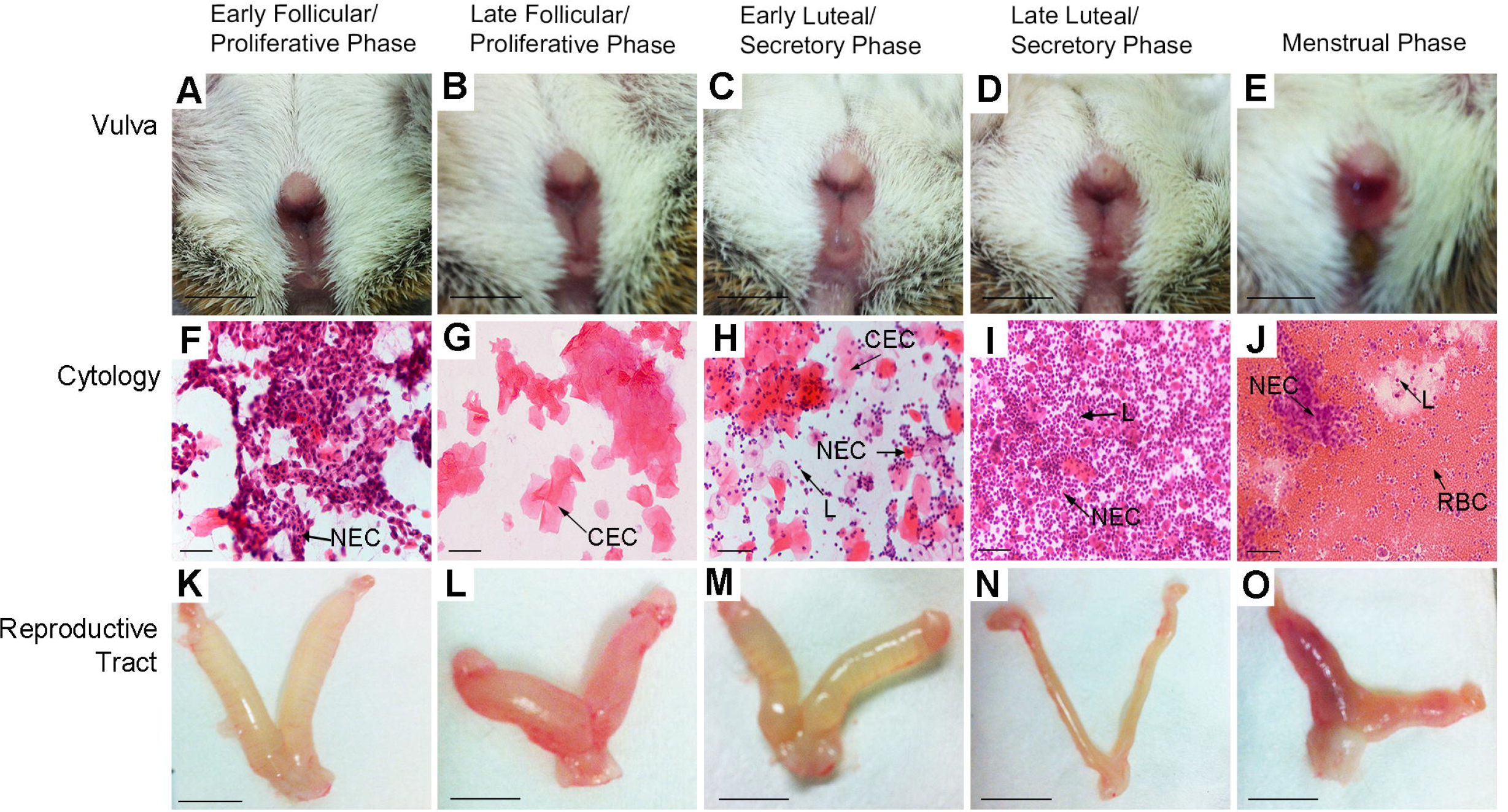
Changes in reproductive tract morphology and cytology across spiny mouse menstrual cycle. <u>Vulva</u> prior to smearing showed a gaping opening in the follicular phase (**A, B**), cellular debris transitioning to the luteal phase (**C**) before closure (**D**). Bloody discharge was observed on the vulva of 29% of females at the time of smearing (**E**). Scale bar = 1 cm. <u>Vaginal cytology</u> (Haematoxylin & Eosin) consisted of nucleated epithelial cells (NEC) (**F**), cornified epithelial cells (CEC) (**G**) and leukocytes (L) (**H, I**). Heavy onset of bleeding during the first day of menses shows high numbers of red blood cells (RBC), some leukocytes and NECs (**J**). Scale bar = 50 μm. <u>Uteri</u> during proliferative phase, were heavily distended with fluid (**K, L**) before resorption (**M**) and complete absence of fluid (**N**) in the secretory phase. Blood was clearly visible in the uterine horns during menses (**O**). Scale bar = 1 cm.

To ensure that bleeding was not caused by vaginal lavage the technique was used in F_1_ (C57BL/6J X CBA/J) hybrid mice. We found no evidence for the presence of red blood cells during any stage of the mouse oestrous cycle (Figure S1) and therefore concluded that the cyclical changes and uterine bleeding seen in the spiny mouse vaginal cytology were a natural phenomenon and not caused by trauma from the lavage technique.

Mallory’s trichrome stain was used to visualise histological changes in the reproductive tract (Figure 3). During the follicular phase of the spiny mouse cycle, the endometrial layer of the uterus was thin (~65 μm). The ovaries contained antral follicles, before rupture, and development into corpora lutea. During the luteal phase, vascular remodelling and decidualisation were observed, endometrial thickness increased 4-5 fold (~320 μm) and the diameter of the uterine lumen decreased. Degeneration of the corpus luteum followed, coinciding with shedding of the endometrium (Table 1 and Figure 3).

**Figure 3:**
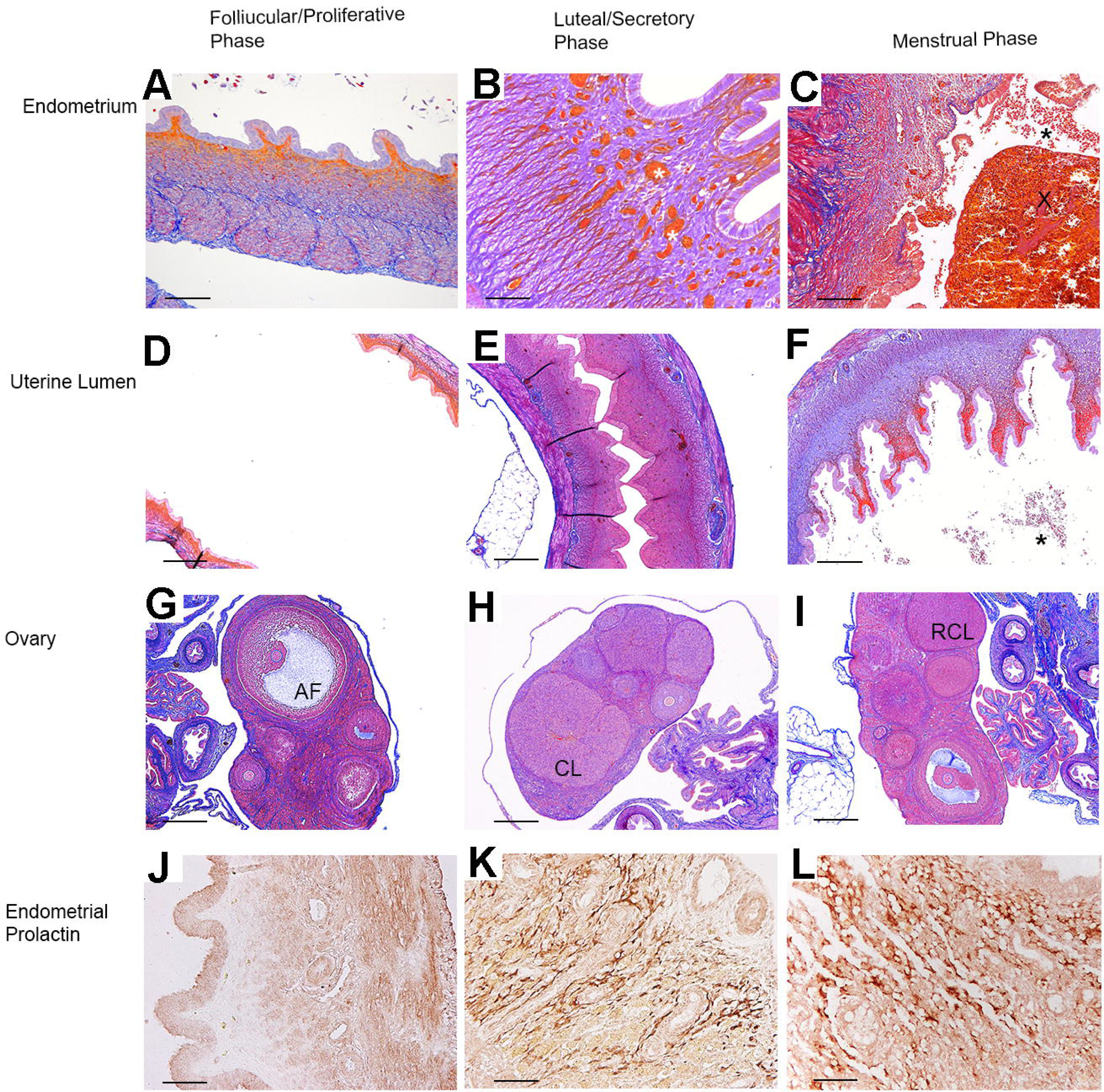
Histological changes of the spiny mouse reproductive tract across the menstrual cycle. Structural changes to the endometrium (scale bar 100μm), uterine lumen (scale bar 200μm) and ovary (scale bar 200μm) are shown. The endometrium during proliferative phase (**A**) is thin before increasing in the secretory phase (**B**), with substantial blood vessel remodelling (white asterisks). During menses (**C**), the endometrium is shed, visible in the peeling of the uterine epithelium, detachment of masses of old endometrial tissue (x) and red blood cell infiltration in the uterine lumen (**F**) (black asterisks). The uterine lumen is wide in diameter during the proliferative phase (**D**) and significantly reduced during the secretory phase (**E**). Antral follicles (AF) are prominent in the ovaries during the follicular phase (**G**) before collapse into corpora lutea (CL) (**H**) and regression (**I**). Immunopositive endometrial prolactin is absent during the proliferative phase (**J**), but secreted from decidualised stromal cells during the luteal phase surrounding newly formed blood vessels (**K**) before the decidual cells are shed during menses (**L**).

Decidualisation of the endometrium was confirmed with immunohistochemical staining for prolactin [17, 20]. Endometrial prolactin staining was absent during ovulation. Immunopositive decidual cells in the endometrium were abundant during the luteal phase (Figure 3). To quantify prolactin immunostaining, we measured optical density. The strongest staining in the endometrium was during the luteal phase, which was followed approximately 2 days later by shedding of the uterine lining.

Plasma progesterone was measured with an ELISA kit (Table S1A and S1B) to determine if histological and immunohistochemical findings were correlated with hormonal changes consistent with the process of menstruation. Significant increases in plasma progesterone concentrations (peaking at 102.1 ng/mL) were noted in the luteal phase compared to all other stages (Table 1). The onset of menstruation-specific cytological and haemorrhagic changes commenced at the time plasma progesterone concentrations begin to fall, a hallmark of spontaneous decidualisation[1, 21]

## DISCUSSION

The spiny mouse reproductive cycle is divided into uterine and ovarian phases consistent with other menstruating species (Figure 4), although this is the first report of such observations in a rodent. Recent hypotheses suggest that menstruation is a consequence of spontaneous decidualisation, and that spontaneous decidualisation evolved in those species with highly invasive fetal tissues and a high rate of embryonic chromosomal abnormality[1, 8]. Why and how menstruation evolved in the common spiny mouse is unknown, but spiny mice share many reproductive features with other menstruating mammals suggesting parallel evolution of these traits with menstruation; for example spontaneous decidualisation, hemochorial placentation, spontaneous ovulation and few, well-developed offspring [16, 22].

**Figure 4:**
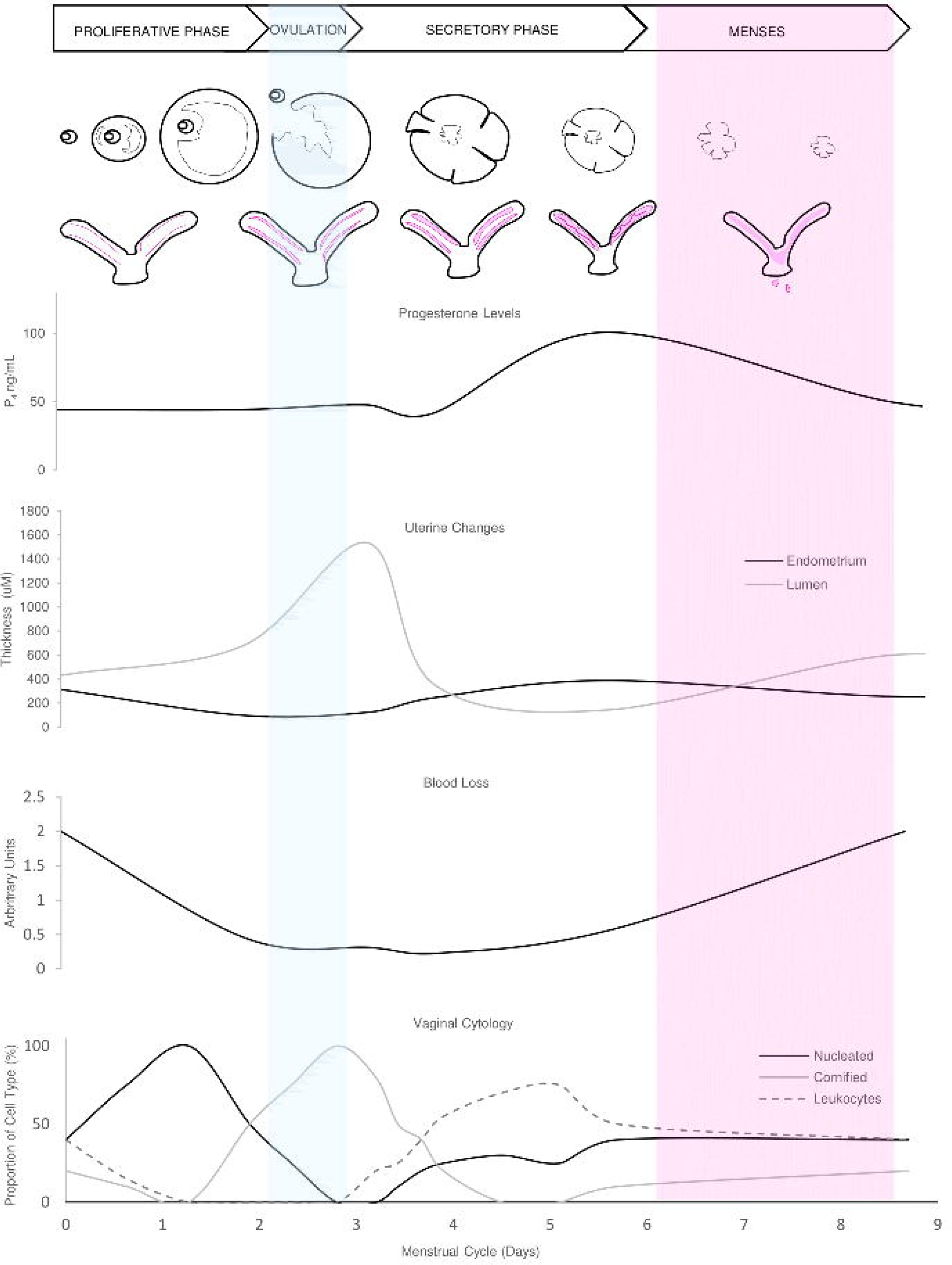
Summary of the spiny mouse menstrual cycle. Cyclical shedding of the endometrium correlates with the regression of the corpus luteum and falling progesterone levels.

The unprecedented discovery of menstruation in a rodent species suggests that this phenomenon was overlooked by previous investigators[16] because of the current dogma that rodents are not menstruating mammals. Studies are now required to determine if menstruation occurs in other species of spiny mice and in the related genera of *Uranomys, Deomys and Lophuromys* and perhaps *Meriones, Gerbillus* and *Tatera*[23–25]. The peripheral position of the *Acomyinae* in most rodent phylogenies[23, 26] suggests that it may be useful and appropriate, in the light of these unusual characteristics of spiny mice, to broaden phylogenetic analysis of this and related genera.

Previous studies have established the spiny mouse as a more useful laboratory species for studies of various aspects of human reproduction and neonatal development than other rodents. The hormonal profile and precocial offspring greatly enhance the utility of this species as a research tool[27, 28]. The spiny mouse should also now be regarded as the most accessible, and cost-effective laboratory model for research into menstruation and menstrual disorders of women, including premenstrual tension (PMT) and endometriosis[29].

## ACKNOWLEDGEMENTS

We acknowledge the efforts and technical assistance of the Hudson Institute of Medical Research histology facility and Monash Health Translational Precinct. We thank Lois Salamonsen and Nikeh Shariatian, for resource allocation and training; Nadia Hale and Lesley Wiadrowski for providing histological and immunohistochemical support; Tim Moss for editing of manuscript.

## SUPPLEMENTARY INFORMATION

### FIGURE LEGENDS

**Figure S1: Vaginal cytology of the female F1 mouse at X200 magnification**. (**a**) Early follicular phase; transition from proestrus to oestrus, containing both nucleated and cornified epithelial cells. (**b**) Oestrus characterised by cornifed epithelial cells. (**c**) Metestrus; transition to luteal phase. (**d**) Early diestrus with large leukocytic infiltration. (**e**) Transition to new fertile cycle. No red blood cells are apparent during any stage of the mouse oestrous cycle. Scale bar = 50 μm.

### TABLES

**Table S1A:** Progesterone spike and recovery results of spiny mouse plasma

| Sample (n) | Level of Analyte Spike | Expected | Observed | Recovery % |
| --- | --- | --- | --- | --- |
| Plasma (4) | Low | 16.7 | 18.8 | 112.5 |
|  | Med | 22.9 | 20.9 | 91.1 |
|  | High | 70.4 | 103.7 | 147.4 |
| *Mean Recovery* |  |  |  | *117.0* |

Recovery of >80% indicates sufficient detection of progesterone in spiny mouse plasma to consider derived values valid for analysis.

**Table S1B:** Progesterone linearity-of-dilution results for spiny mouse plasma

| Sample | Dilution | Expected (Neat) | Observed x Dilution Factor | Recovery % |
| --- | --- | --- | --- | --- |
|  | Neat |  | 26.6 |  |
| High P<sub>4</sub> | 1/4 | 26.6 | 46.6 | 174.8 |
| Plasma | 1/8 |  | 69.5 | 261.1 |
|  | 1/16 |  | 28.9 | 108.3 |

## REFERENCES

1. Emera, D., R. Romero, and G. Wagner, The evolution of menstruation: A new model for genetic assimilation. Bioessays, 2012. 34(1): p. 26–35.

2. Rasweiler, J.J., Spontaneous decidual reactions and menstruation in the black mastiff bat, Molossus ater. The American Journal of Anatomy, 1991. 191: p. 1–22.

3. Brosens, J.J., et al., A role for menstruation in preconditioning the uterus for successful pregnancy. American journal of obstetrics and gynecology, 2009. 200(6): p. 615. e1–615. e6.

4. O’Neil, D. Old World Monkeys. Primates 2014 [cited 2015 12 December 2015]; Available from: http://anthro.palomar.edu/primate/prim_6.htm.

5. The IUCNRed List of Threatened Species. 2015 [cited 2015 09 December]; Available from: www.iucnredlist.org.

6. Demir, R., A. Yaba, and B. Huppertz, Vasculogenesis and angiogenesis in the endometrium during menstrual cycle and implantation. Acta histochemica, 2010. 112(3): p. 203–214.

7. Gellersen, B. and J.J. Brosens, Cyclic Decidualization of the Human Endometrium in Reproductive Health and Failure. Endocrine Reviews, 2014. 35(6): p. 851–905.

8. Clarke, J., The adaptive significance of menstruation: The meaning of menstruation in the elimination of abnormal embryos. Human Reproduction, 1994. 9(7): p. 1204–1207.

9. Zhu, J.-Y., Z.-J. Pang, and Y.-h. Yu, Regulation of Trophoblast Invasion: The Role of Matrix Metalloproteinases. Reviews in Obstetrics and Gynecology, 2012. 5(3-4): p. e137–e143.

10. Huppertz, B., Maternal and fetal factors and placentation: implications for preeclampsia. Pregnancy Hypertension: An International Journal of Women’s Cardiovascular Health, 2014. 4(3): p. 244.

11. Khong, T., The pathology of placenta accreta, a worldwide epidemic. Journal of clinical pathology, 2008. 61(12): p. 1243–1246.

12. Rudolph, M., et al., Induction of overt menstruation in intact mice. PLoS ONE, 2012. 7(3).

13. Brasted, M., et al., Mimicking the events of menstruation in the murine uterus. Biology of Reproduction, 2003. 69(4): p. 1273–1280.

14. Dickinson, H. and D. Walker, Managing a colony of spiny mice (Acomys cahirinus) for perinatal research. Australian and New Zealand Council for the Care of Animals in Research and Training (ANZCCART) News, 2007. 20(1): p. 4–11.

15. Byers, S.L., et al., Mouse Estrous Cycle Identification Tool and Images. PLoS ONE, 2012. 7(4): p. e35538.

16. Peitz, B., The Oestrous Cycle of the Spiny Mouse (Acomys cahirinus). Journal of Reproduction and Fertility, 1981. 61: p. 453–459.

17. Evans, J., et al., Defective Soil for a Fertile Seed? Altered Endometrial Development is Detrimental to Pregnancy Success. PLoS ONE, 2012. 7(12).

18. Monice, F.L., et al., Granulated Decidual Cells in the Mouse Deciduoma: A putative Source of Decidual Prolactin in Mice. Cells Tissues Organs, 2001. 168: p. 252–263.

19. Atik, A., Effects of high-dose caffeine on the cerebrum of the immaute ovine brain, inAnatomy and Developmental Biology. 2014, Monash University: Australia.

20. Handwerger, S., R.G. Richards, and E. Markoff, The Physiology of Decidual Prolactin and other Decidual Protein Hormones. Trends in Endocrinology and Metabolism, 1992. 3(3): p. 91–95.

21. Salamonsen, L. A., Tissue injury and repair in the female human reproductive tract. Reproduction, 2003. 125: p. 301–311.

22. O’Connell, B.A., et al., Sexually dimorphic placental development throughout gestation in the spiny mouse (Acomys cahirinus). Placenta, 2013. 34(2): p. 119–26.

23. Dubois, J.-Y.F., F.M. Catzeflis, and J.J. Beintema, The phylogenetic position of “Acomyinae”(Rodentia, Mammalia) as sister group of a Murinae+ Gerbillinae clade: Evidence from the nuclear ribonuclease gene. Molecular phylogenetics and evolution, 1999. 13(1): p. 181–192.

24. Jansa, S.A. and M. Weksler, Phylogeny of muroid rodents: relationships within and among major lineages as determined by IRBP gene sequences. Molecular phylogenetics and evolution, 2004. 31(1): p. 256–276.

25. Agulnik, S.I. and L.M. Silver, The Cairo spiny mouse Acomys cahirinus shows a strong affinity to the Mongolian gerbil Meriones unguiculatus. Molecular biology and evolution, 1996. 13(1): p. 3–6.

26. Steppan, S.J., et al., Multigene phylogeny of the Old World mice, Murinae, reveals distinct geographic lineages and the declining utility of mitochondrial genes compared to nuclear genes. Molecular phylogenetics and evolution, 2005. 37(2): p. 370–388.

27. Quinn, T.A., et al., Ontogeny of the Adrenal Gland in the Spiny Mouse, With Particular Reference to Production of the Steroids Cortisol and Dehydroepiandrosterone. Endocrinology, 2013. 154(3): p. 1190–1201.

28. Lamers, W.H., et al., Hormones in perinatal rat and spiny mouse: relation to altricial and precocial timing of birth. American Journal of Physiology, Endocrinology and Metabolism, 1986. 251(1): p. E78–E85.

29. Treloar, S.A., et al., Early menstrual characteristics associated with subsequent diagnosis of endometriosis. American journal of obstetrics and gynecology, 2010. 202(6): p. 534. e1–534. e6.

